# Personalized Deep Learning based Source Imaging Framework Improves the Imaging of Epileptic Sources from MEG Interictal Spikes

**DOI:** 10.1101/2022.11.13.516312

**Authors:** Rui Sun, Wenbo Zhang, Anto Bagić, Bin He

## Abstract

Electromagnetic source imaging (ESI) has been widely used to image brain activities for research and clinical applications from MEG and EEG. It is a challenging task due to the ill-posedness of the problem and the complexity of modeling the underlying brain dynamics. Deep learning has gained attention in the ESI field for its ability to model complex distributions and has successfully demonstrated improved imaging performance for ESI. In this work, we investigated the capability of imaging epileptic sources from MEG interictal spikes using deep learning-based source imaging framework (DeepSIF). A generic DeepSIF model was first trained with a generic head model using a template MRI. A fine-tuning procedure was proposed to introduce personalized head model information into the neural network for a personalized DeepSIF model. Two models were evaluated and compared in extensive computer simulations. The MEG-DeepSIF approach was further rigorously validated for imaging epileptogenic regions from interictal spike recordings in focal epilepsy patients. We demonstrated that DeepSIF can be successfully applied to MEG recordings and the additional fine-tuning step for personalized DeepSIF can alleviate the impact of head model variations and further improve the performance significantly. In a cohort of 29 drug-resistant focal epilepsy patients, the personalized DeepSIF model provided a sublobar concordance of 93%, sublobar sensitivity of 77% and specificity of 99%, respectively. When compared to the seizure-onset-zone defined by intracranial recordings, the localization error is 15.78 ± 5.54 mm; and when compared with resection volume in seizure free patients, the spatial dispersion is 8.19 ± 8.14 mm. DeepSIF enables an accurate and robust imaging of spatiotemporal brain dynamics from MEG recordings, suggesting its unique value to neuroscience research and clinical applications.

## Introduction

Electromagnetic Source Imaging (ESI) is the process of estimating the underlying brain electrical activity from noninvasive measurements such as electroencephalography and magnetoencephalography (E/MEG). It has been used to study the sensory systems [1,2], attention [3,4], functional networks [5–7] in both healthy subjects and patients. ESI also plays a substantial role in aiding the diagnosis and surgical planning for drug-resistant focal epilepsy. As one of the major neurological diseases, epilepsy is affecting more than 65 million people worldwide [8], one third of which is drug resistant epilepsy. Brain surgery with the goal to remove the epileptogenic tissue becomes a viable treatment option for these patients, if source localization can be performed accurately. ESI is commonly used to analyze the evolution of brain states or the connectivity patterns during the interictal and ictal periods to identify the epileptogenic tissue [9–14]. Thus, a robust and accurate ESI method is of great importance to both the basic research of brain functions and dysfunctions, and the clinical management of neurological diseases [15,16].

However, ESI is a challenging ill-posed problem since the number of measurements are usually limited to at most a few hundred channels, which are much smaller than the possible source locations in the brain. Theoretically, different cortical activations may generate the same scalp patterns. Thus, conventional ESI methods would use *a priori* constraints as regularization terms to limit the solution space and find a unique solution [17]. ESI then can be formulated as an optimization problem, that is, solving for the brain source distributions that can generate the recorded E/MEG while satisfying the chosen constraints. Various types of regularizations have been proposed and are constantly evolving over the years. Minimum norm estimation and its variations [18,19] or beamforming and scanning approaches [20,21] are widely adopted due to their simple formulations and fast computation. Recent developments have incorporated more details into the prior terms, such as imposing time-frequency constrains [22,23], utilizing spatial adjacency matrix for the dipole distributions [24,25], building hierarchical structures with latent variables [26,27], etc. Albeit these efforts on more realistic modeling of the priors, it remains to be challenging to choose and formulate the correct regularization priors that fully model the complex brain sources and networks. Over-simplified or improper priors would negatively affect the ESI performance since the optimization problem could be inaccurately formulated [28].

Deep learning (DL)-based ESI approaches alleviate the challenges of current ESI approaches. They aim at capturing the correct mapping relationship between signal and source spaces through learning from a large amount of training data, instead of explicitly speculating the features of underlying sources’ distributions. The weights of the interconnected units in a neural network are updated iteratively to minimize the difference between the predicted source and the ground truth in the training data [29], and the statistics and characteristics of underlying sources are implicitly embedded into the network through this process. Deep learning-based source imaging framework (DeepSIF) [30] is recently proposed to provide spatiotemporal imaging of cortical sources from EEG recordings. After the model is trained based on a single head model built from a template magnetic resonance image (MRI), it can still reliably image physiological or pathophysiological brain activities across different subjects and databases.

Although EEG and MEG share ample similarities as they both measure the electromagnetic signals originating from the neuronal currents in the brain [31], they differ in signal properties and the forward process from multiple aspects. For instance, MEG has the merits of measuring the magnetic fields with little impact of the intermediate tissues, especially the low-conductivity skull, with simplified forward modeling of MEG as compared to EEG [32]. On the other hand, EEG electrodes can be directly placed on the scalp while the locations of MEG sensors are fixed, and further away from the brain for conventional SQUID-based systems. Accurate head size and co-registration of the MEG and MRI become more prominent factors for MEG source imaging [33,34]. Moreover, the preference of MEG over tangential current sources means brain regions in sulcal walls will have a stronger representation in MEG over sources on the gyri, producing unique topographical patterns compared to EEG signals. Considering the wide application of MEG technique [35–37] and the differences between the two modalities, it is of interest to rigorously evaluate DeepSIF on MEG recordings, to see if the same forward modeling and training pipeline remains to be effective for MEG source imaging.

In this work, DeepSIF was adapted for MEG ESI where the forward process was calculated for the MEG signal based on a template MRI. Computer simulations were performed to evaluate the performance of this generic DeepSIF (GDeepSIF) model and the impact of head model discrepancies between the training and testing data. Then, we proposed to use the weights of the trained GDeepSIF model as the starting values and continue training the neural network with data from a personalized head model generated based on co-registered individual MRI and channel locations for a personalized DeepSIF model (PDeepSIF). In other words, we used a fine-tuning technique to acquire a PDeepSIF model. Last, we validated the GDeepSIF and PDeepSIF models by comparing their imaging results of interictal spikes to epileptogenic regions including intracranial EEG defined seizure onset zones (SOZ) and resection volumes in a cohort of 29 drug-resistant focal epilepsy patients from two clinical centers. We demonstrated that DeepSIF can return accurate imaging results from MEG recordings, consistent with the clinical findings in epilepsy patients. The additional fine-tuning step for PDeepSIF can alleviate the impact of head model discrepancy and further improve imaging performance significantly.

## Methods

### DeepSIF Outline

The deep learning-based source imaging framework (DeepSIF) is a spatiotemporal ESI method where a deep neural network (DNN) is trained with a synthetic dataset generated by spatial-temporal brain network models. For the spatial model, the cortical surface is segmented into multiple regions and the spatial variations are introduced by a region-based growing method. The source locations are randomly selected among the cortical segments and the size of the source is determined by randomly grouping the neighboring segments with the center segment. The temporal model of each cortical segment is represented by one neural mass model (NMM). NMM describes the average excitation and inhibition behaviors of neuron populations and can generate various types of brain dynamics such as alpha oscillation, evoked potentials, or interictal spikes [38–40]. Using the brain network model, signals with various spatial and temporal patterns can be generated and projected to the scalp based on the anatomical head model. The scalp-brain signal pairs are used as the input-output for the DNN during training. The DNN consists of a spatial module to process the spatial distribution of the scalp data distorted by noise and volume conduction, and a temporal module to model the dynamics of the sources over time and provide the final spatiotemporal estimation of the source distribution (Fig. 1, top half).

**Figure 1.**
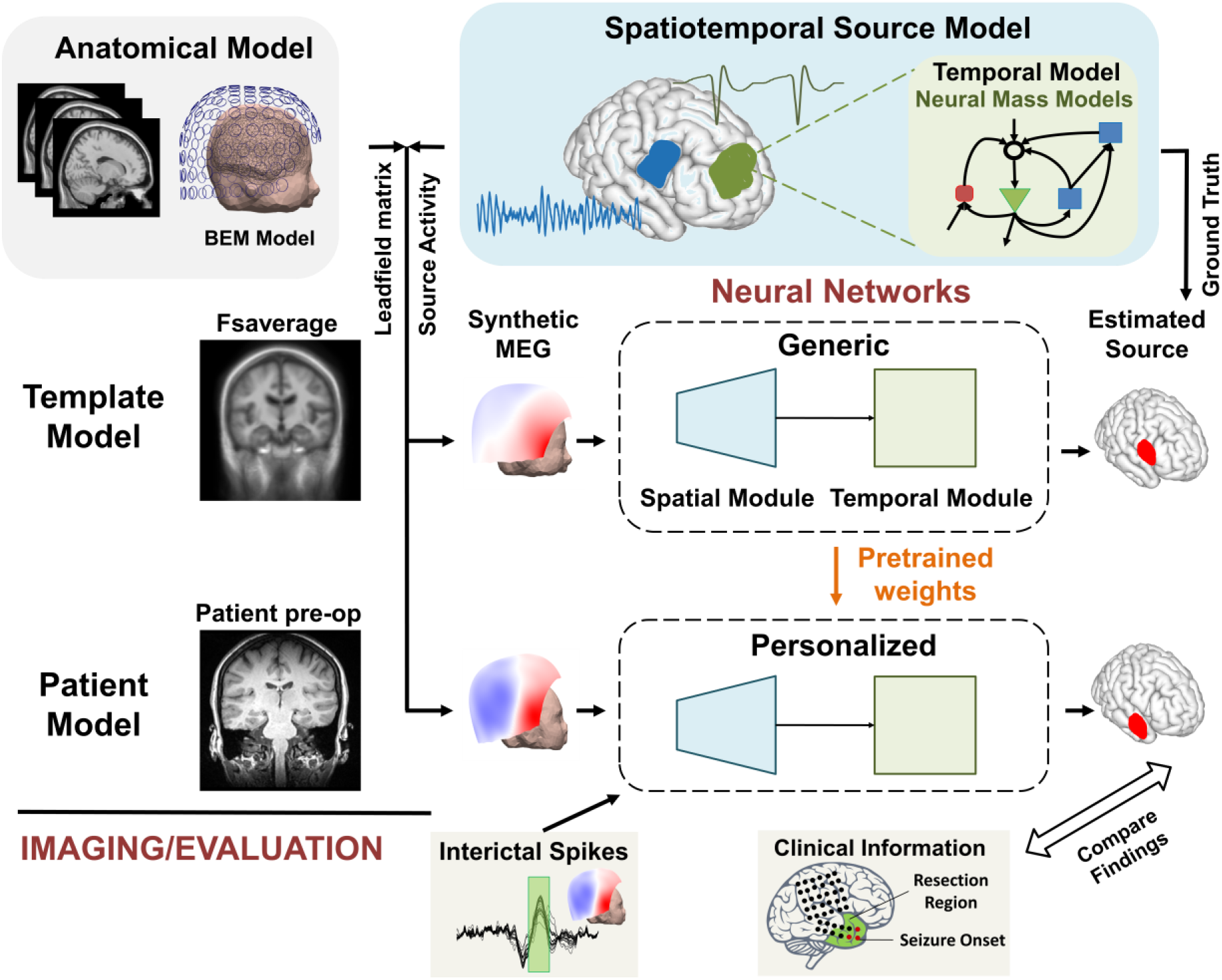
Schematic diagram of study design. The source activities are generated by the spatiotemporal source model, consisting of interconnected neural mass models. A generic head model using template MRI is used to generate synthetic MEG data to train a generic network model. Personalized training data are generated using individual MRI and co-registered MEG sensor locations, which are further used to fine-tune the weights of the generic DeepSIF for a personalized network model.

### Generic DeepSIF Model Training and Evaluation

To generate a generic training dataset for the GDeepSIF model in the simulation study, the template MRI (fsaverage5) [41] was used as the genetic head model with the cortical surface, the skull, and the scalp extracted. The cortical surface was segmented into 994 regions and the Jansen-Rit model [42,43] was used as the NMM for each segment. Then, source patches with different sizes, shapes, locations and temporal waveforms were generated following the same source data generation process as summarized in the DeepSIF outline and detailed in [30]. A 148 MEG magnetometer helmet was used as the MEG sensor configuration. The helmet location was manually adjusted to a centered position and the leadfield matrix was calculated using boundary element method (BEM) model with openMEEG [44] in Brainstorm [45]. The NMM signal was then scaled [30] and projected to the scalp with added Gaussian white noise so that signal to noise ratios (SNR) of 5, 10, 15, or 20 dB were obtained between the scalp signal and noise. In total, a two-source training dataset containing 620,256 spatiotemporal brain-scalp signal samples was generated based on the superposition principle of the two sources.

The network architecture is similar to the one used in [30]. It consisted of a spatial module to pre-filter the MEG signal and a temporal module to model the temporal dynamics. The spatial module had a residual network (ResNet) architecture [46], composed of fully connected layers, which processed the spatial information at each time point independently. Two layers formed a ResNet block (ResBlock) with a skip connection, and Exponential Linear Unit (ELU) activation function [47] was used. For the ResBlock with different input and output dimensions, the skip connection was another fully connected layer to account for the dimension change. The first ResBlock had an input size of 148 (the number of MEG sensors, depending on the MEG systems used. In real patient analysis, the input size was 102 for UPMC MEG data.) and the same output size. The number of MEG sensors could be different across different MEG systems, here we use a 4D Neuroimaging system with 148 magnetometers to perform all the simulation analysis, and we believe the simulation results should generalize to other systems with different numbers of magnetometers. The second ResBlock projected the dimension from 148 to 500, and was followed by another fully connected layer with ELU activation function and output dimension 994. The temporal module aggregated the output from the spatial module over time, and provided the spatiotemporal activity of the source. It had 3 hidden layers and employed Long Short Term Memory (LSTM) [48] with hyperbolic tangent (TanH) activation units. All LSTM layers had an input size of 994 and an output size of 994. Both the source and sensor space signals were scaled by their maximum absolute value to have a maximum or minimum of 1 or −1. During training, the loss function was the mean square error loss (MSE) between the model output and the ground truth source activity. Adam optimizer [49] was used for the training with a weight decay of 1e-6. The learning rate was 3e-4 and the batch size was 64. The whole network was implemented in PyTorch and trained on one NVIDIA Tesla V100 GPU [50].

The source patches in the test dataset were separately generated following the same protocol as the training data. Three test datasets were created using different head models and leadfield matrices to evaluate GDeepSIF’s generalizability. Each dataset contains single source data with 47,712 samples at 5-20 dB SNR levels. First, the same leadfield matrix used in the generic train dataset was adopted for the test dataset when projecting the source space signal to the sensor space. This dataset is called the “no change” dataset. In the second test dataset which is called the “tilt helmet” dataset, the MEG helmet was tilted left for 5 degrees from the original position to simulate the MRI-MEG co-registration error. The leadfield was recalculated using the template MRI and tilted MEG sensor locations. Third, three different subjects’ MRIs were used to evaluate the impact of head geometry. MEG helmets were co-registered to each individual head surface and the leadfield matrices were calculated using subjects’ BEM models. Each leadfield matrix generated one-third (randomly selected) of the test samples in this “different BEM” dataset. Samples in these three datasets were used as the input for the trained GDeepSIF model, and the results are shown in Fig. 2c.

**Figure 2.**
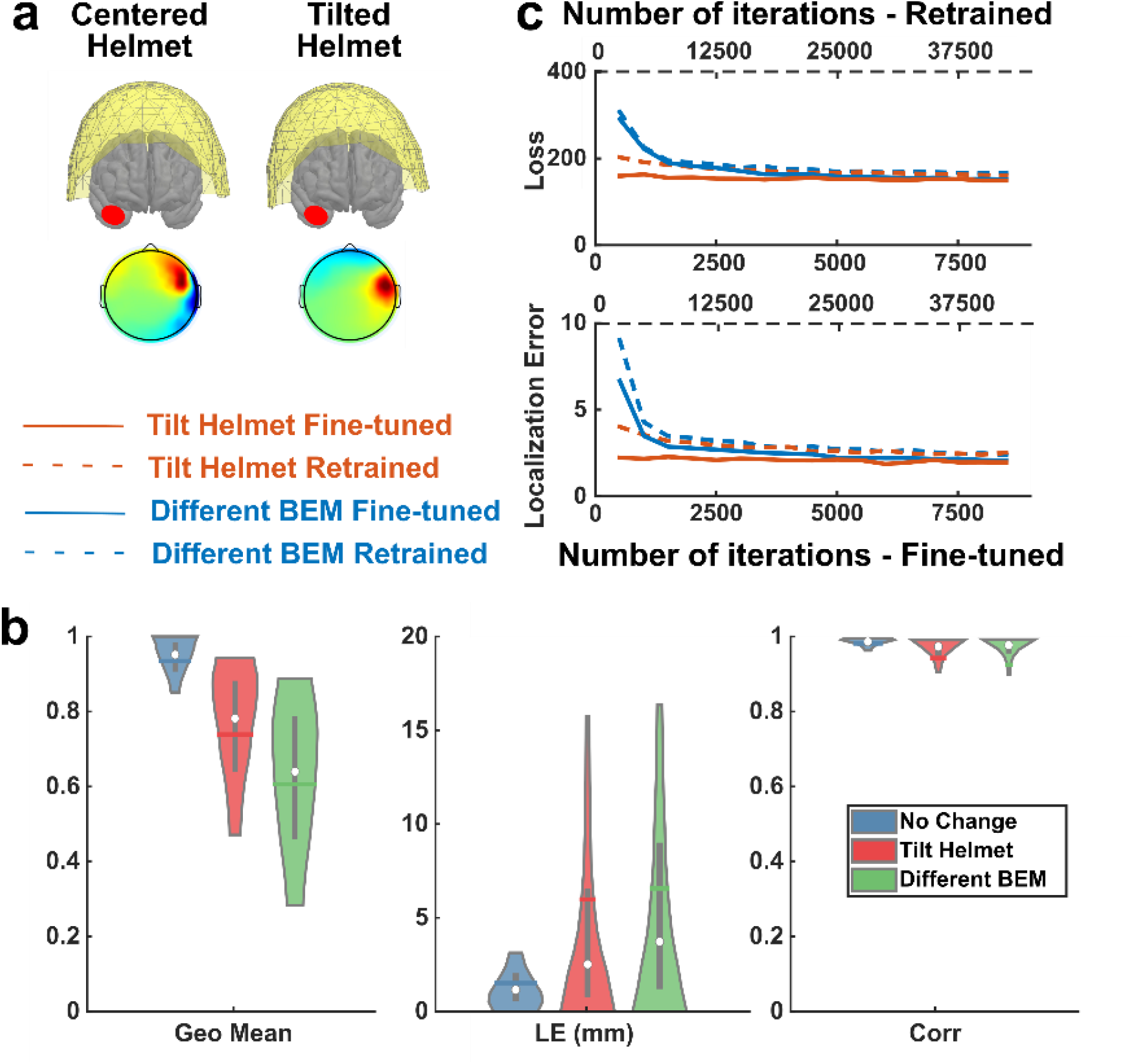
**a**, Example topographical patterns of the same cortical source from two test datasets. **b**, The performance of the GDeepSIF model on three test datasets. The geo mean represents the geometric mean of sensitivity and specificity. Corr represents the linear correlation between the estimated time activity and the simulated time profiles. The dataset names refer to the head model used during the forward process. The distributions are demarcated within the 10th to 90th percentile. The gray bars span the 25th to 75th percentile, the white circle is the median and the colored horizontal bar is the mean of the distribution. (n = 47,712, Geo Mean = 0.93±0.07, 0.74±0.19, 0.61±0.24; LE = 1.52±1.74, 5.98±9.75, 6.57±8.18 mm; Corr = 0.98±0.02, 0.94±0.13, O.93±O.21; for “no change”, “tilt helmet” and “different BEM” respectively). **c**, Loss function and localization error for four different DeepSIF models. Personalized training data were generated by individual head models, either with a tilted helmet or different MRIs (different BEM). DeepSIF trained with “tilt helmet” training data from scratch with random initialization is called tilt helmet retrained (dash line), and continuing training based on GDeepSIF weights is called “tilt helmet finetuned” (solid line). The naming follows the same rule for the “different BEM” dataset.

### Personalized DeepSIF Model Training and Evaluation

The leadfield matrices used in the “tilt helmet” and “different BEM” were also used to generate the personalized training data to either re-train or fine-tune the DNN for PDeepSIF models. The source configurations remained to be the same in the personalized training data, but the leadfield matrices were updated to the corresponding matrices when projecting the source activities to the sensor space. To re-train a DeepSIF model, the extract same training procedures were followed as training a GDeepSIF model, while the training data was replaced with the corresponding personalized data (“tilt helmet” or “different BEM). To fine-tune a DeepSIF model, the weights of the GDeepSIF model were used to initialize the PDeepSIF weights. The weights were continued to be updated using the personalized training data with a learning rate of 1e-4.

The PDeepSIF model fine-tuned with the “tilt helmet” training data was used as an example to validate the performance of the proposed fine-tuning procedure. Test datasets containing one to three-sources were generated with the “tilt helmet” leadfield matrix. There are a total of 47,712 samples at 5-20 dB SNR levels in each dataset.

### Evaluation Metrics for Simulation Study

The Otsu’s thresholding technique [51] was used to identify the boundary of the imaging solution. The algorithm finds a threshold to separate the foreground (active brain sources) and background value that minimizes the intra-class variance. The whole cortical space is denoted by *J*. The estimated active source regions can be denoted by *J_e_*, and the simulated active source regions are denoted by *J_s_*. The estimated and simulated nonactive source regions can be denoted by *J_ne_ = J\J_e_* and *J_ns_ = J\J_S_*. The ability of DeepSIF to estimate the extent, location and temporal dynamics of the cortical sources is evaluated in computer simulations using the following metrics:

#### 1. Sensitivity and Specificity

The sensitivity, also called the true positive rate, is defined as the overlap between the estimated and simulated active sources divided by the simulated active sources, which is calculated as ^|*J_e_* ⋂ *J_s_*|^/_|*J_s_*|_, denotes the size of the sources.

The definition of specificity is the overlap between the estimated and simulated nonactive sources divided by the simulated nonactive sources. However, for source imaging applications, the size of nonactive regions is usually much larger than the size of active regions, directly using *J_ns_* as the nonactive “ground truth” to calculate the specificity would bias the results. Because of this imbalance between the active and nonactive regions, even if *J_e_* is overly diffused compared to *J_s_*, there remains a large *J_ne_* and *J_ne_* ⋂ *J_ns_* area. The original definition of specificity will provide numerically high specificity values, failing to evaluate the method’s true ability to identify the boundary of *J_s_*. A more balanced specificity requires the number of active and inactive source regions to be the same. Thus, a modified specificity as proposed in [25] was adopted. A number of |*J_s_*| regions were randomly selected, either among the neighboring regions of *J_s_* (denoted by 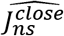) or near the far local maximum of *J_e_* (denoted by 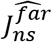). These two types of nonactive regions assess the focality of the estimation and the generation of spurious sources. Here we define the neighboring regions to be within 30 mm Euclidean distance to the source boundary. The final specificity is defined as the average of the *close* and *far* specificity calculated using 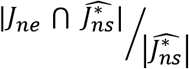, where * represents *close* or *far*. The geometric mean of the sensitivity and specificity can also be calculated as 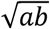, where *a* and *b* represent sensitivity and specificity respectively.

#### 2. Localization errors

For each cortical segment *J_e,i_* in the estimated source *J_e_*, the localization error (LE) is defined as the minimum Euclidean distance to the ground truth regions *J_s_*. One test sample consists of multiple active cortical segments and the LE for one test sample is the mean LE for all segments in the estimated source.

#### 3. Linear correlation

Linear correlation is the Pearson correlation between the solution’s reconstructed waveform and the simulated waveform.

### Patient Information

The clinical data collection were approved by relevant institutions at the University of Pittsburgh Medical Center (UPMC) and Minnesota Epilepsy Group. The data analysis was approved by the Institutional Review Board (IRB) of Carnegie Mellon University. Patients gave their informed consent to participate in this study.

Twenty-nine focal drug-resistant epilepsy patients from these two clinical centers were included in this study (Table 1). All patients underwent a complete presurgical evaluation, including a MEG and intracranial EEG monitoring, as well as a resective surgery. The surgical outcome was scored based on Engel Surgical Outcome Scale [52] by physicians. All patients achieved seizure-free outcome (Engel I) during a follow-up period of at least 12 months. Seven out of 29 patients analyzed were from Minnesota Epilepsy Group. Each patient underwent a 20- to 40-minute MEG session using the Magnes 2500 WH (148 MEG magnetometers, 4D Neuroimaging). The sampling rate is 1,017 Hz. Twenty-two out of 29 patients analyzed were from UPMC. Each patient underwent a 60-minute MEG session using the Elekta Neuromag Vector View 306 Channel System (Elekta Neuromag, Helsinki, Finland). It contains 102 magnetometers and 204 planar gradiometers and only data from the magnetometers were included in the present analysis. The sampling rate is 1,000 Hz.

**Table 1.**

| Patient Information |  |  |  |  |  |
| --- | --- | --- | --- | --- | --- |
| Pt. # | Gender | Surgical Resection | Intracranial EEG | Follow-up Duration | Surgery age |
| 1 | M | Right anterior temporal lobectomy | SEEG over the right frontal temporal region | 2 years | 19 |
| 2 | F | Left anterior temporal lobe lobectomy including mesial structure | SEEG over the left temporal region | 2.5 years | 17 |
| 3 | F | Right anterior temporal lobectomy with amygdalohippocampectomy and resection of seizure focus in right orbital frontal area | SEEG over the right temporal region | 2 years | 42 |
| 4 | F | Right anterior temporal lobectomy including mesial structure | SEEG over the right hemisphere | 3.5 years | 10 |
| 5 | M | Left anterior temporal lobectomy including mesial structures | SEEG over the left frontal temporal region | 1.7 years | 28 |
| 6 | F | right temporal lobectomy and right frontal topectomy | SEEG over the left and right temporal region | 1.5 years | 53 |
| 7 | M | right temporal lobectomy | N.A. | 3 years | 9 |
| 8 | M | Right anterior temporal lobectomy | Right temporal grids | 5 years | 50 |
| 9 | M | Left anterior temporal lobectomy | Left temporal grids and strips | 2 years | 64 |
| 10 | F | A resection in the right supplementary motor area | Right frontal and parietal grids | 3 years | 52 |
| 11 | M | Resection of seizure focus and anterior temporal lobectomy. | Left temporal and frontal grids | 4 years | 43 |
| 12 | M | Right temporal lobectomy | Right temporal grids | 7 year | 25 |
| 13 | F | Left anterior temporal lobectomy | Grids and strips in right frontal, temporal and parietal regions | 3 years | 52 |
| 14 | M | Right temporal lobectomy | Right temporal grids | 7 years | 45 |
| 15 | F | left frontal resection | Grids and strips in left frontal, temporal and parietal regions | 6 years | 30 |
| 16 | M | A right frontal lobe seizure focus resection | Right frontal grids | 6 years | 32 |
| 17 | F | Resection of her left temporal and orbitofrontal epileptogenic zone | Left temporal grids and strips | 4 years | 52 |
| 18 | F | Left temporal lobectomy | SEEG over the left temporal region | 4 years | 42 |
| 19 | F | Right temporal lobectomy | SEEG over the right temporal and frontal region | 3 years | 30 |
| 20 | M | Left frontal resection | SEEG over the left and right frontal region | 1 year | 46 |
| 21 | F | Removal of the right frontal AVM | SEEG over the right parietal region | 3 years | 17 |
| 22 | F | Left temporal lobe resection | SEEG over the left temporal region | 5 years | 27 |
| 23 | F | Resection of the presumed transmantle cortical dysplasia in the right frontal lobe. | SEEG over the right temporal and frontal region | 3 years | 44 |
| 24 | M | Left transfusiform amygdalohippocampectomy | SEEG over the left and right temporal region | 4 years | 54 |
| 25 | F | right anterior temporal lobectomy with hippocampal and amygdala resection | SEEG over the right temporal region | 2 years | 60 |
| 26 | M | left anterior temporal lobectomy | SEEG over the left temporal and parietal region | 5 years | 36 |
| 27 | F | left temporal lobectomy | Grids and strips in left frontal, temporal and parietal regions | 4 years | 50 |
| 28 | M | resection | SEEG over the right temporal and occipital region | 7 years | N.A. |
| 29 | M | resection of a left temporal lesion | SEEG over the left and right temporal region | 1 year | 20 |

### Clinical MEG Data Analysis

A GDeepSIF model was trained for each clinical center, as the number of MEG channels, thus, the input size of the neural network, is different (148 for Minnesota Epilepsy Group and 102 for UPMC). The PDeepSIF model for each subject was acquired by finetuning the GDeepSIF model. For each subject, the cortical surface, the skull, and the scalp were extracted, and its cortical space was segmented following the same segmentation atlas as used in the generic brain model in Freesurfer [41]. The surfaces were then imported to Brainstorm and the cortical surface was down-sampled to 20,487 vertices using iso2mesh [53]. The alignment of the anatomical and functional fiducial points was performed manually, and then the helmet positions were further refined based on the digitalized head points and scalp surface. The leadfield matrix was calculated using BEM for each patient, which was used to generate the personalized training data to fine-tune the DeepSIF model.

A 10-minute segment of the MEG recording was bandpass filtered between 1 and 40 Hz and down-sampled to 500 Hz. Interictal spikes were extracted, averaged and scaled by the maximum absolute value of the data to range −1 and 1 before being used as the input for the trained DeepSIF model. Three conventional source imaging methods were used as the benchmark methods. The imaging results for standardized low resolution brain electromagnetic tomography (sLORETA) [19] and linearly constrained minimum variance (LCMV) beamformer [20] were calculated using MNE-Python (version 0.22.0) [54]. Coherent maximum entropy on the mean (CMEM) [25] was calculated using the brainentropy plugin (version 2.7.3) in Brainstorm. The output source reconstruction was averaged for a 20 ms window around the peak of the spike and the Otsu’s method was used to find the extent of the imaging solution when evaluating the performance of all methods [51].

In patient studies, the sensitivity, specificity and spatial dispersion (SD) were calculated with respect to the resection region. The definitions of sensitivity and specificity are the same as the simulation study and the resection region is used as the ground truth (or active source regions). Spatial dispersion is defined as the weighted mean of the distance of each reconstructed region to the resection area. 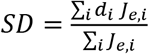, where *d_i_* is the minimum distance to the resection region for reconstructed region *i*, and *J_e_* is the estimated source map. When comparing to the SOZ electrodes, sublobar concordance, sublobar sensitivity and specificity, and localization errors were calculated. The cortical surface was segmented into 20 subregions (Anterior temporal, lateral posterior temporal, medial posterior temporal, orbitofrontal, prefrontal, premotor, central, superior parietal, inferior parietal and occipital). The imaging result is concordant with SOZ if the maximum of the source estimate and the SOZ electrodes are located in the same subregions. When calculating the sublobar sensitivity and specificity, the subregions containing both source estimates and SOZ electrodes are defined as the true positive subregion, and the true negative subregions are defined as regions containing neither SOZ nor source estimates. The sublobar sensitivity definition is the same as the sensitivity defined above, which is the number of true positive subregions divided by the number of SOZ subregions. The sublobar specificity is calculated as the number of true negative regions divided by the number of subregions without SOZ. Note that unlike the specificity comparing to the resection, all the nonactive subregions were used for the evaluation without sampling. When evaluating the localization error, two LEs are calculated. The first LE is defined as the average distance of each intracranial EEG defined SOZ to the closest estimated source regions. The second LE is defined as the average distance of each estimated source region to the closest SOZ. SOZ LE is defined as the average value of these two LEs (Fig. 4a).

**Figure 3.**
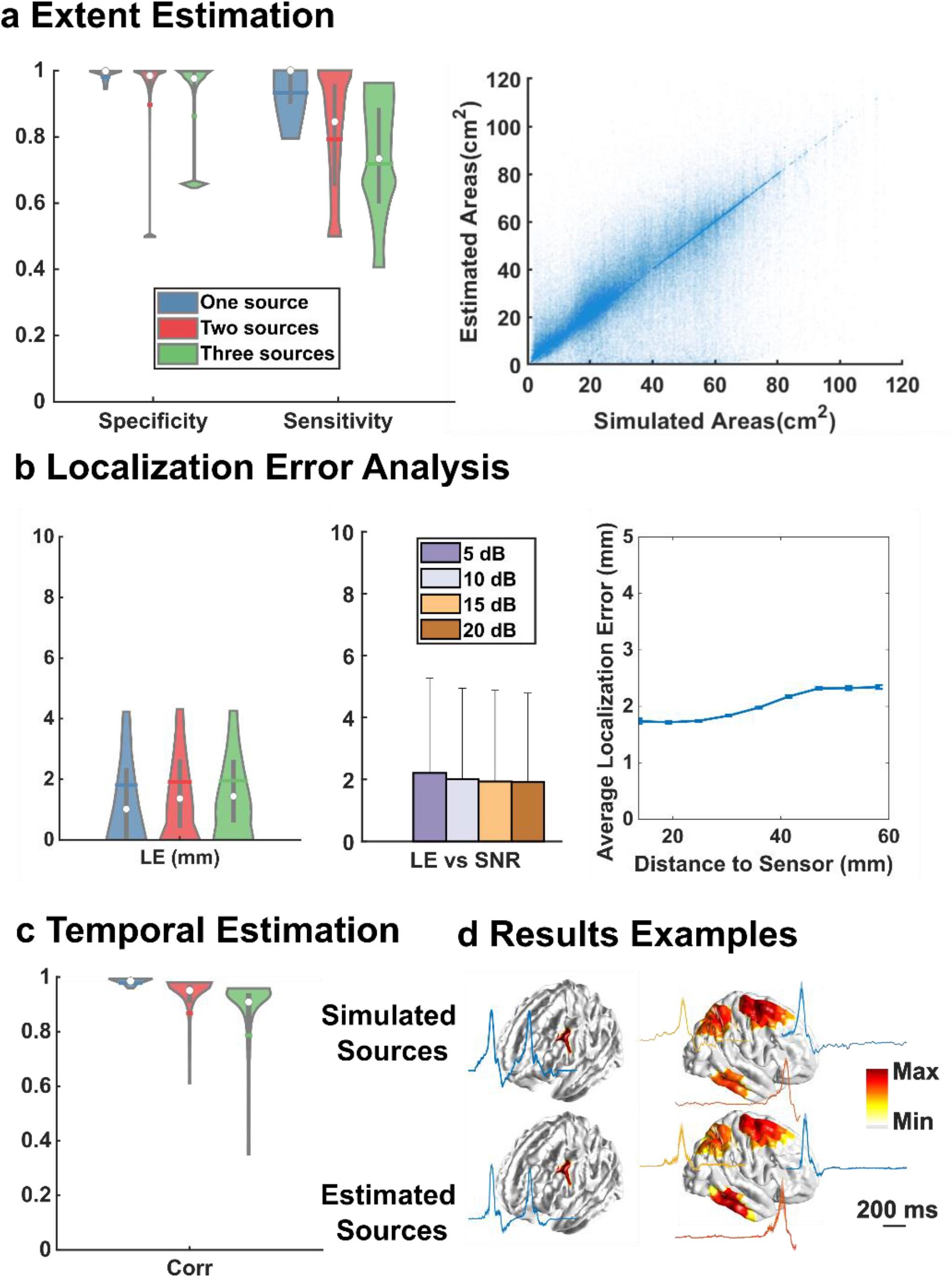
Model performance on the personalized test datasets. **a**, Extent estimation. Left: The specificity (n = 47,712, 1-source: 0.98±0.04, 2-sources: 0.90±0.18, 3-sources: 0.86±0.20) and sensitivity (n = 47,712, 1-source: 0.93±0.11, 2-sources: 0.78±0.21, 3-sources: 0.71±0.20) of three test datasets with one-, two-, and three-sources, respectively. Right: Simulated source area vs estimated source area for three datasets combined. **b**, Localization error analysis. Left: the LE distributions of each dataset (n = 47,712,1-source: 1.81±3.01, 2-sources: 1.91±2.21, 3-sources: 1.96±2.09 mm). Middle: LE vs SNR for all three datasets combined, the error bar shows the standard deviation. Right: LE vs depth, the plot shows the average LE for all sources within a particular depth, and the error bar shows the standard error of the mean. **c**, Temporal estimation. The linear correlation between ground truth and reconstruction of each dataset (n = 47,712, 1-source: 0.98±0.03, 2-sources: 0.87±0.21, 3-sources: 0.79±0.25 mm). **d**, Imaging examples. Source locations and waveforms of ground truth and reconstructed activities for a single source (left) and three sources (right).

**Figure 4.**
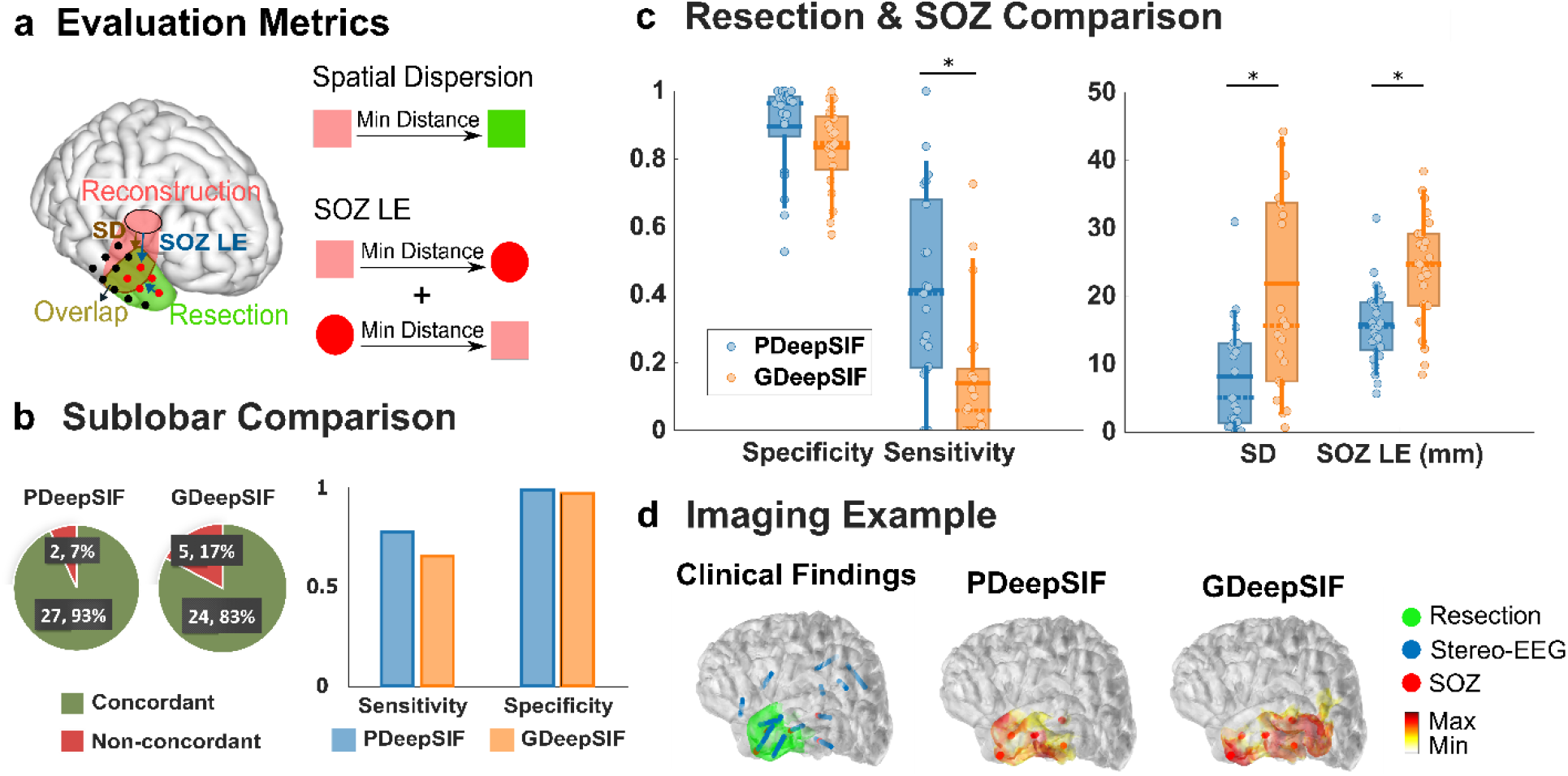
Clinical validation for GDeepSIF and PDeepSIF models. [P] and [G] represent PDeepSIF and GPDeepSIF respectively. **a**, Evaluation metric illustration (details in methods). **b**, Sublobar comparison results. Left: concordance rate (93% [P], 83%[G]). Right: Sublobar sensitivity (77% [P], 66% [G]) and specificity (99% [P], 97% [G]). **c**, Quantitative spike-imaging results. The horizontal solid line shows the mean, the dashed line shows the median, the bars span the 25th to 75th percentile of the data, the vertical bars span the 10th to 90th percentile of the data, and each circle represents individual patients. (Specificity: n = 21, 0.90 ± 0.14 [P], 0.83 ± 0.12 [G]; Sensitivity: n = 21, 0.41 ± 0.29 [P], 0.14 ± 0.20 [G]; Spatial dispersion (SD): n = 21, 8.19 ± 8.14 mm [P], 21.90 ± 19.03 mm [G]; SOZ LE: n = 27, 15.78 ± 5.54 mm [P], 24.86 ± 10.40 mm [G].) **d,** Examples of spike-imaging results along with the surgical resection outcome and iEEG defined SOZ.

## Results

### GDeepSIF Performance in Computer Simulations

Fig. 2b shows the GDeepSIF model performance evaluated on three test datasets. When there is no change between the head model used for the train and test datasets, GDeepSIF model can achieve excellent localization performance with a LE of 1.52 ± 1.74 mm. However, when the head model used is modified in the test dataset, the accuracy decreased by an expected however noticeable amount. For the “tilt helmet” test dataset, the LE becomes 5.98 ± 9.75 mm, and for the “different BEM” test dataset, the LE is 6.57 ± 8.18 mm. This is mostly caused by the change in the scalp-brain relationship as shown in Fig. 2a. For the same cortical source, the tilted helmet measures a different topographical pattern (right) compared to the original training data (left), which caused a shift of the estimated source regions and a decreased sensitivity and specificity value.

### PDeepSIF Training and Evaluation

To further improve the performance on the personalized test datasets, DNN can be retrained from random initial weights or finetuned based on a trained GDeepSIF model using personalized training data. The training curves in Fig. 2c followed a similar trend, however, the number of iterations to fine-tune a model (solid line, bottom axis), is only one fifth of retraining a model (dash line, top axis). Fine-tuning GDeepSIF model provided faster convergence compared to retraining the network. The number of iterations means the number of DNN weights updates, which roughly translates to the training time. With one fifth of the training time, fine-tuning a DNN can reach the same validation loss as retraining the network from scratch. When evaluating the changes of LE during training, a similar trend can also be observed that the LE converged to a low value with fewer updates when fine-tuning based on GDeepSIF weights.

The fine-tuned DeepSIF model was further evaluated for its ability to estimate the extent (Fig. 3a), location (Fig. 3b) and temporal dynamic (Fig. 3c) of the underlying source activities on the personalized test dataset. PDeepSIF can estimate the temporal evolution of the source activities with a high correlation value (0.88 ± 0.17). High specificity (0.91 ± 0.17) and sensitivity (0.81 ± 0.20) are also achieved when averaged across the one-, two-, and three-source conditions for all SNR levels. A significant Pearson’s correlation of 0.83 is achieved (p < 0.001) between the simulated source areas and estimated source areas, which means PDeepSIF can identify the extent of the cortical sources with high accuracy. The average localization error is 1.89 ± 2.47 mm for all source configurations across datasets. When examining the samples at different SNR, the LEs are 2.06 ± 2.47, 1.88 ± 2.44, 1.82 ± 2.52 and 1.80 ± 2.39 mm for SNR = 5, 10, 15, 20, respectively. Source depth is defined as the distance between the source center to the closest MEG sensor. Robust performance can be observed when the source depth increases as the change of the average LE is less than 1 mm. PDeepSIF can also provide consistent superior performance when the inter-source distance or the temporal correlation between sources varies (Supplementary Fig. S1) in the two-source configuration, indicating PDeepSIF’s capability in distinguishing multiple closely located or correlated sources. The simulation study demonstrated that PDeepSIF can provide accurate and robust source estimates from MEG measurements.

### Comparison of GDeepSIF and PDeepSIF in Patient Data Analysis

We compared the interictal spike imaging results of GDeepSIF and PDeepSIF in a cohort of 29 focal epilepsy patients. All patients suffered from drug-resistant epilepsy and went through intracranial EEG (iEEG) monitoring and resective surgery with seizure free outcomes. Twenty-one out of 29 have clear post-operative MRI images to extract the resection regions, and 27 out of 29 patients have post-implantation computer tomography (CT) images available to identify the iEEG electrode locations.

Both GDeepSIF and PDeepSIF models can produce reasonable results close to the clinical findings, achieving a high sublobar concordance rate of 83% and 93% respectively. However, the PDeepSIF performed better in all metrics compared to GDeepSIF (significant in resection sensitivity, SD and SOZ LE, two-sided Wilcoxon signed rank test, p<0.01), demonstrating that for MEG DeepSIF, finetuning is a critical step to improve the imaging accuracy. As we can see in one imaging example in Fig. 4d, the GDeepSIF model introduced a bias in the estimate, away from the clinical ground truth, to compensate for the discrepancies in the leadfield matrix. After the finetuning, PDeepSIF can produce results with a better concordance with the resection region. This can also be observed in the group-level metrics (Fig. 4c). Although GDeepSIF produced results with high spatial specificity (Sublobar specificity equals 97%, resection specificity equals 83%), demonstrating that GDeepSIF can still provide an good extent estimate without sources at spurious locations, a “shift” can be observed for GDeepSIF because of the inaccuracies in the training leadfield matrix, as reflected in the increased SD and SOZ LE values. However, the reconstruction remained to be in close proximity to the clinical ground truth as the mean SOZ LE is 24.86 mm. On the other hand, after the model is finetuned with personalized head model information, the SOZ LE dropped to 15.78 mm, indicating excellent concordance with the SOZ covered area.

### Comparison of PDeepSIF and Other Methods in Patient Data Analysis

Fig. 5 shows the comparison of PDeepSIF to other benchmark ESI methods: LCMV, sLORETA and CMEM. PDeepSIF demonstrated superior imaging performance, with an SD value of 8.19 ± 8.14 mm and SOZ LE of 15.78 ± 5.53 mm, which is significantly better when compared to the benchmark methods. Although LCMV and sLORETA can provide high sensitivity, their low specificity impairs their ability to identify the true epileptic regions. PDeepSIF can provide a reasonable extent of the sources, without being too focal or overly diffused as it has a balanced sensitivity and specificity value. When calculating the geometric mean (GM) of the sensitivity and specificity, PDeepSIF reaches 0.54 ± 0.28, and it is significantly better than 0.43 ± 0.23, 0.41 ± 0.21 and 0.30 ± 0.30 for LCMV, sLORETA and CMEM respectively (two-sided Wilcoxon signed rank test, p<0.05). These validation results demonstrated that PDeepSIF can reliably image and localize the epileptogenic tissue from MEG measurements.

**Figure 5.**
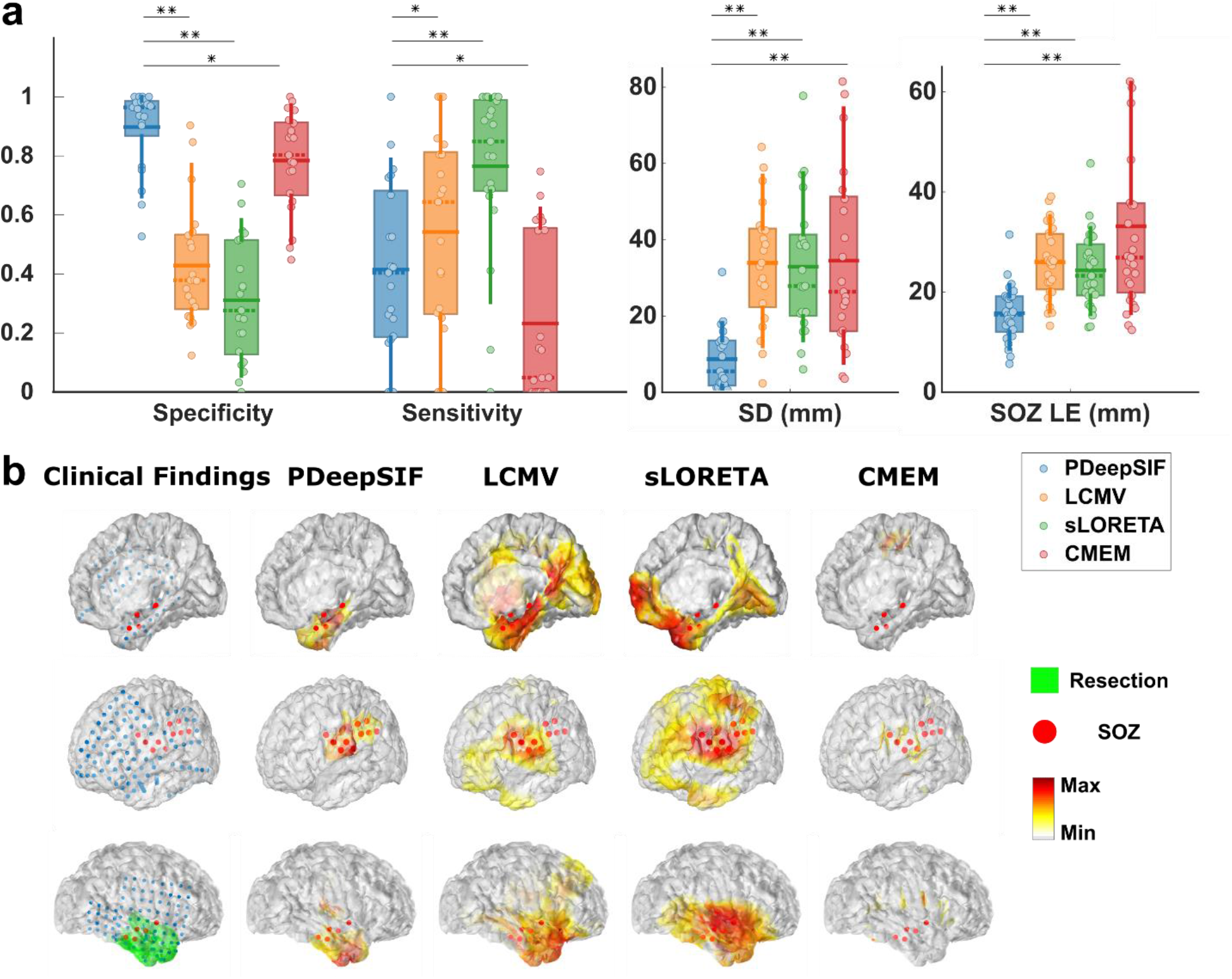
**a**, Quantitative interictal spike imaging results. (Specificity: n = 21, 0.90 ± 0.14 [P], 0.43 ± 0.20 [L], 0.31 ± 0.21 [S], 0.78 ± 0.17 [C]; Sensitivity: n = 21, 0.41 ± 0.29 [P], 0.54 ± 0.34 [L], 0.76 ± 0.28 [S], 0.23 ± 0.28 [C]; Spatial dispersion (SD): n = 21, 8.19 ± 8.14 [P], 33.36 ± 16.16 [L], 32.36 ± 17.94 [S], 33.95 ± 23.60 [C] mm; SOZ LE: n = 27, 15.78 ± 5.53 [P], 25.96 ± 7.06 [L], 24.34 ± 7.52 [S], 33.14 ± 18.42 [C] mm, where [P], [L], [S] and [C] represent PDeepSIF, LCMV, sLORETA and CMEM respectively). Paired two-sided Wilcoxon signed rank test was used with statistical significance cutoffs of (*P<0.05, **P<0.01). **b**, Examples of spike-imaging results along with the surgical resection outcome and iEEG defined SOZ.

## Discussion

Imaging spatiotemporal brain activities from MEG measurements has become an indispensable tool for research and clinical applications [16]. We further developed a recently proposed deep learning-based source imaging framework and rigorously validated its performance in computer simulations and in a cohort of 29 drug-resistant focal epilepsy patients. We have demonstrated that DeepSIF can be successfully applied to MEG measurements and provide good imaging results with a generic head model. Personalized anatomical information can be incorporated into the framework by continuing to train the generic DeepSIF model with personalized training data. The fine-tuned model can significantly improve the overall performance for estimating the location, extent, and temporal activities of the brain sources across different subjects. The validation of DeepSIF on MEG measurements is a crucial advancement for DL-based electromagnetic source imaging (ESI) methods.

Designing and optimizing the appropriate priors for the ill-posed ESI problem has been a challenging task. Deep learning methods have the advantage of implicitly learning the source distributions instead of explicitly formulating the regularization terms, providing opportunities for a more accurate and robust ESI estimate. There have been several attempts recently to image brain activities using deep neural networks [55–60]. They have shown excellent performance in computer simulations, demonstrating the power of DL-based ESI methods. However, since synthetic training data is required due to the lack of real recordings [61], it is of great importance to evaluate the generalizability of the trained network on a group of subjects with different anatomical features and signal properties. A recent study [30] filled this gap by validating the trained DeepSIF network on a group of epilepsy patients and showed that it can provide accurate and robust imaging results in real EEG recordings. However, most of these studies, including DeepSIF, were developed and validated under the EEG setup. Although EEG and MEG share ample similarities, and many ESI methods can be applied to both simply by changing the leadfield matrix in the forward problem to the corresponding modality, rigorous validation is required before applying DL-based ESI methods on MEG measurements, as the differences in the experimental setup, forward processes, signal properties, etc, could make the training process designed for EEG signals inappropriate for MEG measurements.

In this work, we demonstrated, in systematic simulations, that DeepSIF could be successfully adapted to process MEG measurements, reaching a median LE of around 4 mm for the GDeepSIF model even when a different head model was used in the test data. After fine-tuning the GDeepSIF with personalized training data, the median LE decreased to around 1 mm. When evaluating on the interictal spikes from focal epilepsy patients on a sublobar level, both GDeepSIF and PDeepSIF can provide a high sublobar concordance rate (83 % and 93%). Various sublobar concordance rates have been reported for MEG (47% to 91 %) [62–64] and EEG interictal spike imaging (36% to 95%) [10,65]. In terms of sublobar sensitivity and specificity, there is a tradeoff between the two metrics depending on the properties of the source imaging methods evaluated. For more diffused imaging methods, a sublobar sensitivity of 84%-98% and specificity of 38%-60% can be reached, and the geometric mean of the sensitivity and specificity is in the range of 56% − 77% [66,67]. When using sparse imaging methods for a higher spatial specificity value, the sensitivity could drop to 25%-40% [62]. Correctly estimating the extent of the sources objectively remains to be a challenging problem [11,68]. DeepSIF provides a reasonable balance between identifying the correct source regions and having spurious activations, as the geometric mean of the sublobar sensitivity and specificity is 87% and 78% respectively for PDeepSIF and GDeepSIF. Both GDeepSIF and PDeepSIF models can reach a high specificity (> 97%) without excessive sacrificing on the sensitivity (> 66%). It is observed that GDeepSIF performance is worse compared to PDeepSIF performance due to the variations among the patient head models. However, GDeepSIF could still be used as a valuable and efficient tool for guiding the implantation of iEEG electrodes on a sublobar level, as it has shown high concordance with the SOZ subregions.

The differences between the GDeepSIF model and PDeepSIF model were more significant when compared to the clinical ground truth quantitatively. The spatial specificity compared to the resection of GDeepSIF remains to be above 0.8, however, the sensitivity has a huge decrease. This indicates that, even though GDeepSIF can correctly identify the subregions of the source activity, there is a shift of between the clinical ground truth and the GDeepSIF estimate, reducing the overlap area between the resection and source estimates. This can also be observed in the increased SOZ LE (24.86 ± 10.40 mm). On the other hand, personalized head model information can further reduce the error caused by the shift of the GDeepSIF source estimation, improving the sensitivity value and reducing localization error, as shown in the patient example. The SOZ LE decreased to 15.78 ± 5.53 mm for PDeepSIF, and the SD decreased to 8.88 ± 8.25 mm, demonstrating that excellent source localization results can be obtained in real patients when personalized head models are used.

Using a generic head model (template MRI with standard channel locations) for MEG source imaging is less established than EEG. Studies have shown that when evaluating the effect of head models variations on the leadfield matrices and ESI results, the major differences were caused by the MEG sensor co-registration errors other than the shape of the volume conductor or the measurement noise [34,69]. As SQUID MEG sensors are not directly placed on the scalp, head points describing the head shape as well as the relative positioning between the head and helmet are necessary for an accurate MRI and MEG co-registration and imaging analysis [70]. This posed a unique challenge for GDeepSIF method as the differences between the generic and personalized head model, especially the sensor locations, are not negligible for MEG.

To introduce the correct head shape information to DeepSIF, personalized training data can be used. Training a separate DeepSIF model individually from scratch inevitably increased the computational burden, and we proposed to use the personalized training data to fine-tune the GDeepSIF model to reduce the training time to acquire a personalized model by five times (Fig. 2c). As shown in Fig. 2c, GDeepSIF already learned the mapping relationship between the source and sensor space and can provide reasonable imaging results even for data generated by different head models. Fine-tuning based on GDeepSIF weights can provide a faster convergence and significantly reduce the training time for PDeepSIF model, without sacrificing the imaging performance. We demonstrated that the fine-tuned PDeepSIF can provide accurate and robust source reconstructions under various simulation conditions (multiple simultaneous sources, low SNR, deeply located sources, etc). As shown in Fig. 5, PDeepSIF can also provide accurate imaging results of interictal spikes from epilepsy patients, with high spatial specificity of 90% and low SOZ LE (15.78 ± 5.53 mm), which means the source estimates are concordant with the clinical findings without spurious activations at other locations, an important advantage over other benchmark ESI methods. Note that even if we can achieve a very high specificity value, the sensitivity value compared to resection volume is around 0.4, which means the estimated sources from interictal spikes are within but smaller than the resected region.

Since multiple factors can influence the resection volume in practice, starting with anatomo-electro-clinical correlation, patient’s other clinical characteristics, including individual functional level, identified cognitive deficits, treatment goals (e.g., “cure” vs. meaningful seizure reduction or control/reduction of the most disabling seizures), imaging and functional testing findings, encountered individual anatomical reality during an operation, neurosurgeon’s assessment of risks and even possibly their practice style, the resection region is not necessary the ground truth for the interictal spike activities [71,72]. However, it is still a valuable benchmark to validate our imaging results, especially in seizure free subjects.

In sum, we have demonstrated that DeepSIF can provide robust capability of imaging brain activity from MEG recordings. The present results indicate that even with a generic head model, GDeepSIF can still offer good capability of localizing epileptiform activity at sublobar level. With the additional fine-tuning step with personalized data, PDeepSIF can accurately estimate the extent, location, and temporal dynamics in computer simulations, and provide interictal spike imaging results concordant with clinical findings. This extension of DeepSIF from EEG to MEG is a crucial step for the development of ESI methods in general. However, it is worth noting that, the conclusions were drawn based on the magnetometers of the SQUID MEG systems. Gradiometers have the advantage of reducing the environmental noise during the recording [32], while a higher sensitivity for detecting mesial temporal spikes has been reported for magnetometers [36]. The influence of different sensor configurations on ESI methods remains to be an open question that worth exploring in future investigations [68,73].

## Acknowledgement

This work was supported in part by National Institutes of Health grants NS096761, EB021027, AT009263, MH114233, EB029354, and NS124564, and by a gift from the Pittsburgh Health Data Alliance. R.S. was supported in part by a Fellowship from the Center for Machine Learning and Health. The work used the Extreme Science and Engineering Discovery Environment (XSEDE), which was supported by National Science Foundation grant number ACI1548562. Specifically, it used the Bridges-2 system, which is supported by NSF award number ACI-1928147, at the Pittsburgh Supercomputing Center (PSC).

We thank Dr. Abbas Sohrabpour, Dr. Shuai Ye, and Xiyuan Jiang, for their valuable discussions and assistance in the data analysis. We are grateful to Drs. Alexandra (Popescu) Urban, Gena Ghearing, Arun Anthony, Viji Rajasekaran, Nirav Barot, James F, Castellano, Mark R. Richardson, Luke Henry, and Joseph Mettenburg, for their respective clinical roles in evaluating and treating surgically included patients at UPMC. We also acknowledge selfless dedication and invaluable efforts of the University of Pittsburgh Comprehensive Epilepsy Center (UPCEC) team and particularly the staff of the UPMC Presbyterian University Hospital (PUH) Epilepsy Monitoring Unit (EMU) led by Cheryl Plummer, BS, R. EEG T., NA-CLTM, FASET. We acknowledge the physicians of Minnesota Epilepsy Group, Dr. Meysam Kabrieaei of Children’s Minnesota, and Dr. Kyle Nelson of Allina Health for their clinical roles in evaluating and treating patients. Special thanks to Drs. Deanna Dickens, Nitin Agarwal, Paul Atkinson, and Jessica Winslow for their assistance. We also wish to thank our research coordinator Ms. Sarah Ellis and lead EEG/MEG technologist Mr. Brian Owens for his technical assistance. We thank participating patients and their families whose involvement and sacrifice made this work possible.

## Data availability

The main data supporting the results in this study are available within the paper and in the supplementary materials. Other additional data are available upon reasonable request from the corresponding author.

## Contributions

R.S. and B.H. contributed to developing analysis algorithms and performing data analysis. A.B. and W.Z. contributed to the data collection. B.H. contributed to the supervision of the project. R.S. and B.H. contributed to the initial draft of the manuscript. All authors contributed to the revision of the manuscript.

## Supplementary Information

### Figures

**Figure S1.**
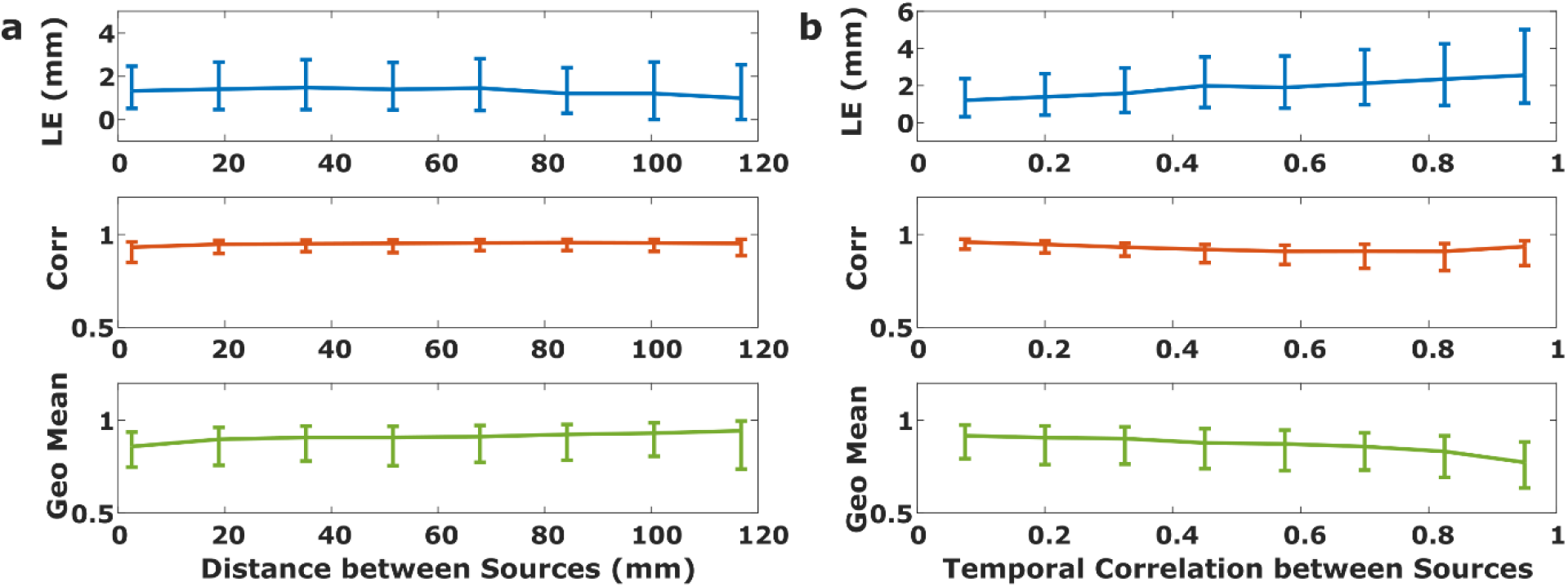
The relationship between the imaging performance and **a**, the distance or **b**, the temporal correlation between sources for the two-source scenario in computer simulations. The plots depict the median localization error, linear correlation over time, and the geometric (geo) mean of sensitivity and specificity (SNR = 5, 10,15, and 20 dB) when the distance or temporal correlation between the two sources is varied. The error bar shows the 25th to 75th percentile. The distance between sources is defined as the minimum distance between the two extended sources.

**Figure S2.**
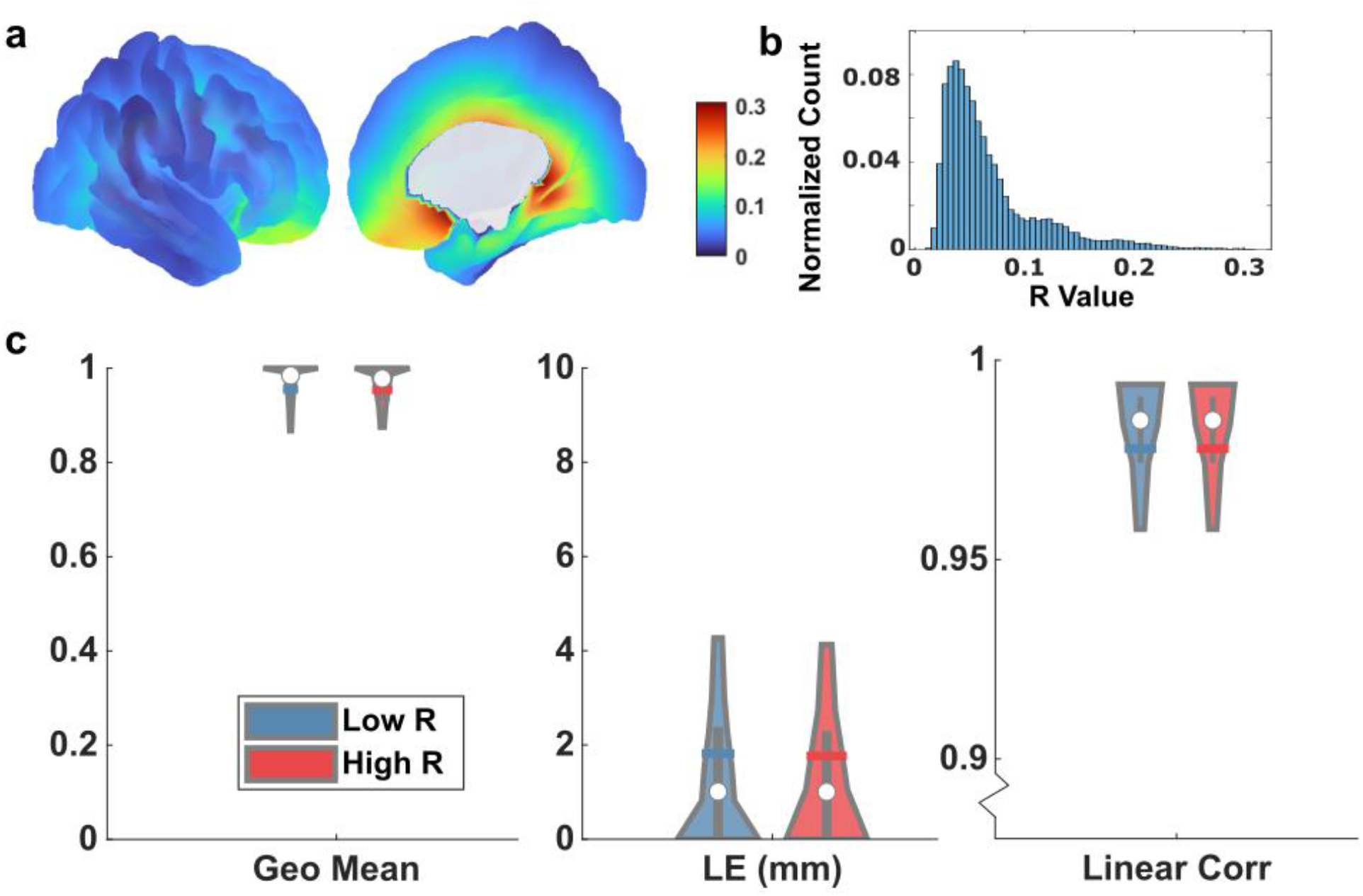
The ratio between minimum and maximum singular value of the lead-field matrix at each source location (Nchannel*3) was defined as the R value to indicate the sensitivity to a certain source orientation. A low R value indicates the difference between the minimum and maximum gain of the leadfield is large, which means that the measurement is more sensitive to a certain orientation. **a,** R value distribution over the cortex. **b,** R value histogram. **c,** PDeepSIF performance for low R value and high R value conditions for a single source dataset. R value for a source patch is calculated as the mean R value of all the vertices in the source patch. Low and high value is divided based on the median of the R distribution (median = 0.07). The geo mean represents the geometric mean of sensitivity and specificity. Corr represents the linear correlation between the estimated time activity and the simulated time profiles. The distributions are demarcated within the 10th to 90th percentile. The gray bars span the 25th to 75th percentile, the white circle is the median and the colored horizontal bar is the mean of the distribution.

**Figure S3.**
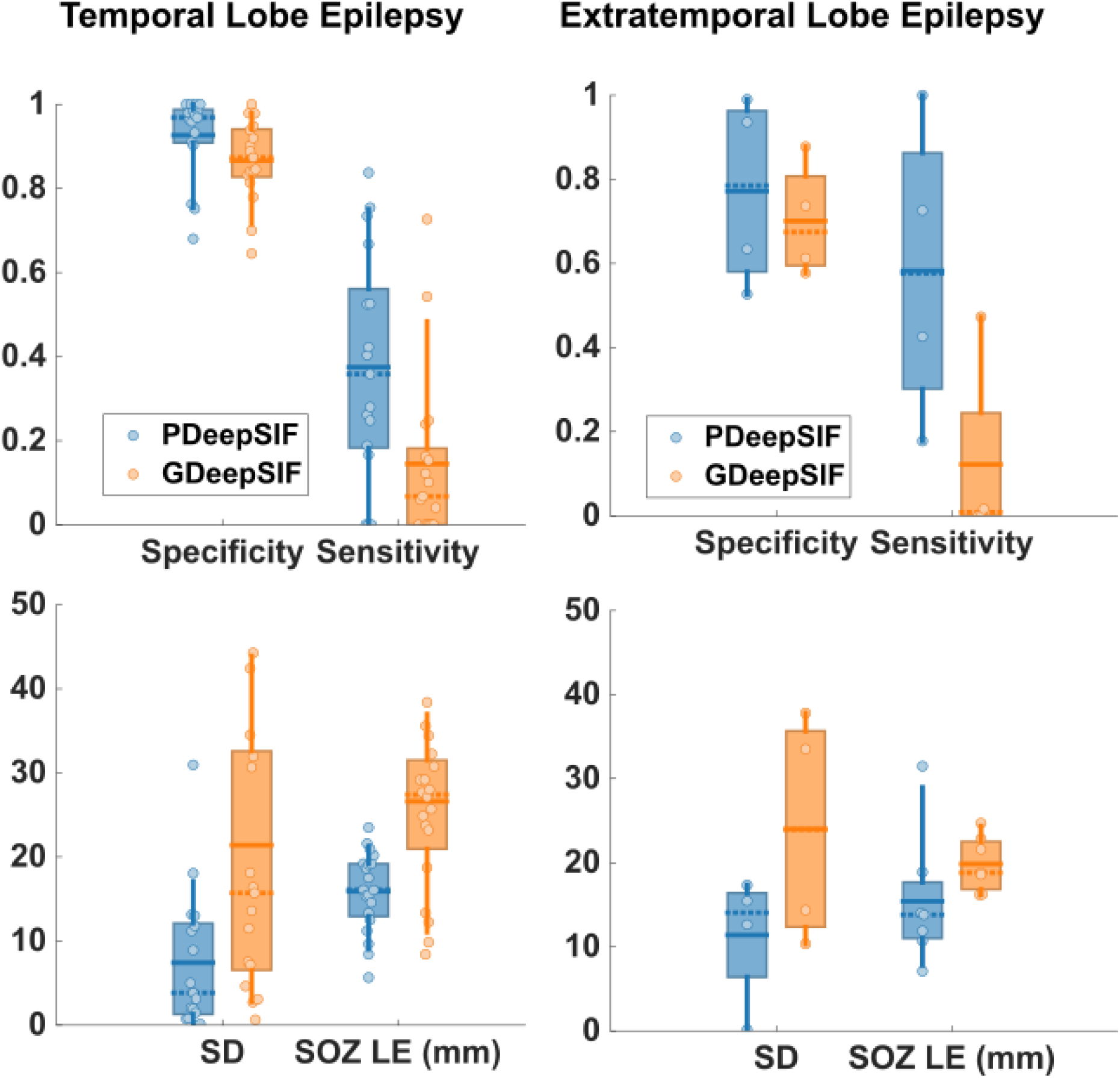
Clinical validation for GDeepSIF and PDeepSIF models for temporal lobe epilepsy (TLE, n=22) and extratemporal lobe epilepsy (ETLE, n=7). [P] and [G] represent PDeepSIF and GPDeepSIF respectively. TLE: Specificity – 0.93 ± 0.10 [P], 0.87 ± 0.10 [G]; Sensitivity – 0.37 ± 0.27 [P], 0.14 ± 0.20 [G]; SD – 7.43 ± 8.27 mm [P], 21.40 ± 20.40 mm [G]; SOZ LE – 15.91 ± 4.69 mm [P], 26.61 ± 11.49 mm [G]. ETLE: Specificity – 0.77 ± 0.23 [P], 0.70 ± 0.14 [G]; Sensitivity – 0.58 ± 0.35 [P], 0.12 ± 0.23 [G]; SD – 11.41 ± 7.74 mm [P], 24.01 ± 13.67 mm [G]; SOZ LE – 15.41 ± 7.94 mm [P], 19.86 ± 3.28 mm [G]. There is no statistical significance between TLE and ETLE for all metrics (Two-sided Wilcoxon rank sum test, p>0.19).

